# Single cell-scale spatial transcriptome profiling of the adult cycling mouse uterus

**DOI:** 10.1101/2025.09.22.677812

**Authors:** Elizabeth Ung, Tyler J. Gibson, LeCaine J. Barker, Elle C. Roberson

**Affiliations:** School of Medicine, Department of Pediatrics, Section of Developmental Biology at the University of Colorado Anschutz Medical Campus

## Abstract

The adult uterus is highly regenerative during the reproductive cycle (menstrual in humans; estrous in mice), while the uterus prepares for a possible pregnancy. Until recently, the regenerative capacity of the mouse uterus was under-appreciated. Therefore, how uterine cell types and tissue compartments coordinate transcriptional and cellular changes across the estrous cycle is poorly understood. To begin to uncover the spatiotemporal molecular regulation of cell remodeling and regeneration, we conducted Visium HD spatial transcriptomics to analyze the adult cycling nulliparous mouse uterus. We integrated transcriptional data binned into 8x8 µm spots across all four cycle stages to annotate cell types. This dataset provides highly resolved transcriptional information of uterine cell types, including luminal epithelium, glandular epithelium, mesenchyme, blood vessels, immune cells, and smooth muscle. These data can be used to define subpopulations of uterine cells and how they change in the adult cycling uterus or determine where disease-related genes are spatially expressed. Collectively, rigorous analysis of this dataset will significantly advance our understanding of uterine biology.

## Background and Summary

The uterus is required for pregnancy, thus is critical for fertility^1^. The inner uterine compartment – the endometrium – is the site of embryo implantation^2^. The outer uterine compartment – the myometrium – synchronously contracts during labor to expel the developed fetus^3^. The ability of the uterus to successfully develop a fetus depends on the health of the non-pregnant uterus, during the menstrual or estrous cycle^2^. During the menstrual cycle, the endometrium sheds during menstruation, then regenerates and remodels the glandular and luminal epithelium, underlying stromal fibroblasts, and vasculature^4,5^. Recent work has established that the mouse endometrium is similarly regenerative during the estrous cycle, where the luminal and glandular epithelium regenerates and the stromal fibroblast population increases by proestrus, and then these same populations simplify or decrease by diestrus^6–8^.

Researchers have used several technologies to explore cellular and molecular characteristics of the uterus. Healthy human uterine samples during the menstrual cycle and labor have been processed for single cell RNA sequencing (scRNAseq)^7,9–13^, single nucleus Assay for Transposase-Accessible Chromatin using sequencing (ATACseq)^14^, and lower resolution spatial transcriptomics^13^. Human uterine samples from patients with intrauterine adhesions^15^, endometriosis^16,17^, recurrent implantation failure^18^, or SARS-CoV-2 infection^19^ have been processed for scRNAseq. The cycling mouse uterus has been probed by scRNAseq and lower resolution spatial transcriptomics^20^, ATACseq^21^, bulk RNAseq^22^, and microarray analysis^23^. The pregnant mouse uterus has been analyzed by scRNAseq at gestational day (GD) 3 and GD4^24^, GD6^25^, and in preterm labor^26^, by bulk RNAseq at GD3 and GD4^27^, and by lower resolution spatial transcriptomics at GD7.5^28^. The mouse uterus has also been profiled by bulk RNAseq following uterine injury^29^ and obesity^30^. These studies have yielded immense understanding of uterine gene expression dynamics following myriad physiological alterations, and the complex cell heterogeneity underlying uterine biology.

However, the listed techniques disrupt tissue architecture as a necessary part of the protocol, limiting our understanding of the spatial organization of these gene expression and cell heterogeneity changes. Moreover, the estrous cycle is often overlooked, so there are few datasets that cover the entire cycle. To overcome these knowledge gaps, we performed unbiased single-cell-scale spatial transcriptomics on the wild type cycling mouse uterus at each stage of the estrous cycle with 10X Visium HD technology^31^. We obtained high quality, spatially resolved, genome-wide transcriptional data from each cycle stage. We use these datasets to identify spatially localized uterine cell types and their gene expression signatures across the estrous cycle. Robust validation of this dataset will significantly advance our understanding of uterine remodeling and regeneration during the estrous cycle, and its role in female fertility. This dataset represents a profound resource to investigate the spatial patterning of uterine cell types and genes required for fertility or disrupted in pathological settings.

## Methods

### Experimental Overview

Our goal was to sequence the single-cell-scale spatial transcriptome of the uterus at each stage of the estrous cycle in sexually mature wild type mice. In brief, we estrous cycle tracked wild type female mice to collect uterine samples at each stage of the estrous cycle, including proestrus, estrus, metestrus, and diestrus (**Figure 1A-B**). We embedded in paraffin (**Figure 1C**), then sectioned each sample with the microtome (**Figure 1D**). After H&E staining according to the 10X Genomics Visium HD protocol (**Figure 1E**), we imaged at 20X (**Figure 1F**), then submitted samples for CytAssist capture, library prep, and sequencing.

**Figure 1.**
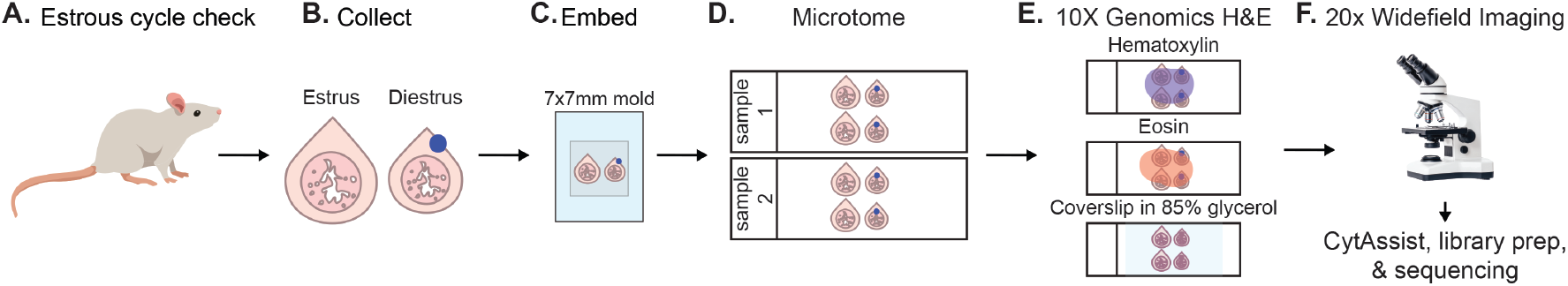
Overview of collected samples and data generation. A) Swiss Webster female mice between 8-10 weeks of age were estrous cycle tracked. B) Uterine samples were collected at each cycle stage (proestrus, estrus, metestrus, and diestrus). Two samples were embedded together (ex: estrus and diestrus) so to differentiate between samples, blue tissue dye was dotted onto one sample per block. C) Uterine samples were paraffin embedded together into a 7x7mm mold. D) Paraffin blocks were sectioned at 5µm using a Leica microtome. E) Sections were hematoxylin and eosin stained following the 10X Visium HD protocol, and then coverslipped in 85% glycerol. F) Sections were imaged at 20X on a Leica DM6b widefield microscope, and then samples were prepped using the 10X CytAssist for subsequent library preparation and sequencing.

### Mice

All experiments were performed on 8 to 10-week-old Swiss Webster female mice that were obtained from Charles River, Inc (USA). Mice were housed in individually ventilated cages in a pathogen-free AALAC-accredited facility with continuous food and water and a controlled 14hr light / 10h dark cycle, in accordance with National Institutes of Health guidelines. All experiments were approved by the University of Colorado Anschutz Institutional Animal Care and Use Committee (IACUC protocol #01267).

### Estrous cycle tracking

Animals were estrous cycle tracked as previously described^23^. In brief, mice were acclimated to estrous cycle tracking for a week by briefly swabbing the vaginal canal using a cotton swab wetted with 1X PBS. Then, animals were tracked by smearing the swab onto a microscope slide, dipping in crystal violet for 45 seconds, rinsing with water, and staging under a Leica DM500 brightfield microscope (Leica Microsystems). Estrous stages were called based on presence and type of epithelial cells or leukocytes^32^.

### Tissue collection and processing

#### Dissections

All dissection tools and dishes were sprayed with RNaseAway (Thermo Fisher Scientific). 8-10-week-old females in the desired estrous stage were humanely euthanized through the isoflurane drop method followed by cervical dislocation. The entire reproductive tract of each animal was harvested and placed into a petri dish filled with chilled 1X PBS/nanopure water. The uterus was bisected at the cervix and extra blood vessels and fat were removed. One uterine horn was placed onto a strip of index card and laid flat before placing into 4% PFA/nanopure water for fixation for 3 hours at 4°C. Samples were then washed in 1X PBS/nanopure water 3x5 minutes, rocking at room temperature.

#### Tissue processing

A 2mm thick slice of fixed uterine tissue was placed into a labelled paraffin cassette. Metestrus or diestrus samples had a dot of blue tissue dye to help distinguish between samples (**Figure 1B**), with the goal of pairing proestrus and metestrus, and estrus and diestrus samples together in a single paraffin block. The samples were incubated with 70% ethanol/nanopure water overnight at room temperature. The next day, samples were placed at 4°C until sent for paraffin processing.

#### Histology processing

The 2mm chunk was sent to the University of Colorado Anschutz Pathology - Research Histology Shared Resource (RRID: SCR_021994) for paraffin embedding. Proestrus and metestrus, and estrus and diestrus samples were paired together, embedding both slices of tissue cut side down in a 7x7mm mold (**Figure 1C**). Embedded samples were stored at 4°C until use.

### Tissue sectioning

All surfaces were wiped down with ethanol, followed by RNaseAway. Paraffin blocks were placed in an ice/nanopure water bath to cool down the paraffin wax before being placed into a Leica microtome. The block was exposed, and tissue was transversely cut at 5µm. Ribbons were placed into a warm nanopure water bath and maneuvered to be placed onto clean microscope slides, aiming for a 15x5 mm pre-drawn rectangle. Another slide was prepared with eight sections to prepare for quality control via fluorescence *in situ* hybridization (RNAscope). Slides were left to dry at room temperature overnight, then stored at 4°C until ready for histology staining or drop off to the University of Colorado Anschutz’s Human Immune Monitoring Shared Resource (HIMSR, RRID: SCR_021985). Additionally, 4 ribbons/block were placed into a pre-chilled, RNase-free tube to perform FFPE RNA extraction (Qiagen RNeasy FFPE Kit, ref #73504) to assess RNA quality.

### RNA quality assessment

We performed three sets of RNA quality control. First, we isolated RNA from ribbons and used a Nanodrop 2000 (Thermo Fisher Scientific) to determine concentration and purity levels. Second, we sent the collected RNA to the Functional Genomics Core at the University of Colorado Anschutz to assess RNA purity using the Tapestation (Agilent). Finally, we sent our sections to the HIMSR to perform RNAscope with positive control probes.

### Hematoxylin and eosin staining

We followed the *Visium HD FFPE Tissue Preparation Handbook* (10x Genomics, CG000684 Rev A) as follows. Bake slides at 60°C for 2 hours. Allow the slides to cool to room temperature. Immerse in xylene 2x, then incubate for 10 minutes. Repeat. Immerse in 100% ethanol 2x, then incubate for 3 minutes. Repeat. Immerse in 96% ethanol 2x, then incubate for 3 minutes. Repeat. Immerse in 70% ethanol 2x, then incubate for 3 minutes. Repeat. Immerse in ultrapure water 2x, then incubate for 20 seconds. Add 1mL of hematoxylin to the slide, incubate at room temperature for 1 minute, and remove. Immerse 5x in ultrapure water. Immerse 15x in fresh ultrapure water. Repeat with fresh ultrapure water. Add 1mL of bluing buffer to the slide, incubate at room temperature for 1 minute, and remove. Immerse 15x in fresh ultrapure water. Immerse slides in alcoholic eosin at room temperature for 1 minute. Immerse the slide in fresh ultrapure water for 30 seconds. Immerse 15x in fresh ultrapure water. Coverslip in 85% glycerol.

### Imaging

Slides were imaged with a 20X objective on a Leica DM6b microscope (Leica Microsystems) with the affiliated LAS-X software. One section from each estrous stage was selected for sequencing based on overall uterine architecture integrity. Slides were sent to the University of Colorado Anschutz’s Genomics Core for further processing.

### Library preparation and sequencing

After imaging, tissue sections underwent pretreatment and were hybridized overnight with 10x Genomics’ whole transcriptome probe pairs according to the *Visium HD Spatial Gene Expression Reagent Kits User Guide* (CG000685 Rev A). Post-hybridization, sections were washed and ligated to form probe pairs. The processed tissue slide and Visium HD slide were then loaded onto the CytAssist instrument to transfer and spatially barcode the ligated probes. Spatially barcoded products were used for library preparation per the Visium HD protocol. Final libraries were pooled and sequenced on an Illumina NovaSeq X Plus system (150 bp × 10 bp × 10 bp × 150 bp) at the University of Colorado Genomics Shared Resource (RRID: 021984), generating approximately 500 million reads per capture area.

### Spatial transcriptomics data analysis

Loupe Browser (version 8.1.2) was used to manually outline tissue sections. Each uterus on the same slide was marked separately, and downstream processing was performed separately. Raw data were processed using Spaceranger (version 3.1.2). Spaceranger outputs data at the level of the 2 μm detection spots present on the Visium HD slide. Due to the sparsity of the 2 μm data, Spaceranger also bins the 2 μm data into larger square bins tiled across the capture area. Unless otherwise noted, downstream analysis was performed using 8 μm square bins, which provided the optimal tradeoff between spatial resolution, quality of cell type identification from clusters, and computational performance. Integration and clustering were performed using Seurat (version 5.1.0)^33^. 8 μm bins with fewer than 50 UMIs were filtered out and data were log normalized. To facilitate efficient clustering of such a large dataset, we used sketch-based sampling to select 50,000 8 μm bins for each stage while preserving rare cell types^34^. This subset was used to perform variable feature selection, data scaling and principal component analysis (PCA). Integration of data from the different stages of the estrous cycle was performed using Seurat’s reciprocal PCA method. 8 μm bins were clustered using graph-based clustering (30 principal components, resolution = 0.8). Integration and clustering were projected onto the full dataset. Differential gene expression was used to identify marker genes for each cluster. *De novo* and known marker genes were used alongside the spatial positions and distributions of each cluster to perform cell type identification. Although 8 μm binning performs better for clustering and cell-type annotation, the 2 μm data provide better spatial resolution for the visualization of individual genes. To reduce the sparsity of the 2 μm data while retaining the high spatial resolution, we utilized SAINSC (version 0.3.1) to perform kernel density estimation (KDE) on the 2 μm data, providing high-resolution, smoothed gene expression profiles for visualization^35^.

### Data visualization

Visualizations of expression data and clusters projected onto the tissue image were generated using SpatialData (version 0.3.0) and Matplotlib (version 3.10.0)^36,37^. Additional graphs were generated with ggplot2 (version 3.5.2)^38^. Plotgardener (version 1.14.0)^39^ was used for assembling figures.

## Data Record and Code Availability

This dataset contains 10X Visium HD spatial dataset consisting of mouse uterus at each estrous cycle stage. Data will be made public upon paper publication in NCBI GEO and will be shared upon request. All analysis code will be made available on GitHub upon paper publication.

## Technical Validation

### Sample quality control and histology

To generate our dataset, we used paraffin embedding to maintain accurate tissue architecture and we co-embedded two different samples per paraffin block. This strategy allowed us to maximize area on the Visium HD slide (10X Genomics, USA). We performed sample quality control to ensure that our workflow resulted in high quality tissue. First, we used a combination of nanodrop and Tapestation to assess our 260/280 ratio and confirmed we had high quality total RNA (**Figure 2A**). As this approach destroys tissue architecture, we also performed RNAscope with control probes (**Figure 2B**). RNAscope showed that our tissue samples had intact RNA and confirmed our workflow produced high RNA quality within the tissue. Low resolution images of the H&E-stained uterine sections were captured with the CytAssist (**Figure 2C**). Because Visium HD provides high resolution transcriptomics data, it was important to also obtain high resolution images of the H&E-stained uterine sections before sequencing (**Figure 2D**). While the original alignment of the transcriptomic data was performed using the low resolution CytAssist images, we wanted to project the transcriptomic data onto the high-resolution images. To accomplish this, we manually detected our tissue in the low-resolution image and aligned that with the high-resolution image (**Figure 2E**).

**Figure 2.**
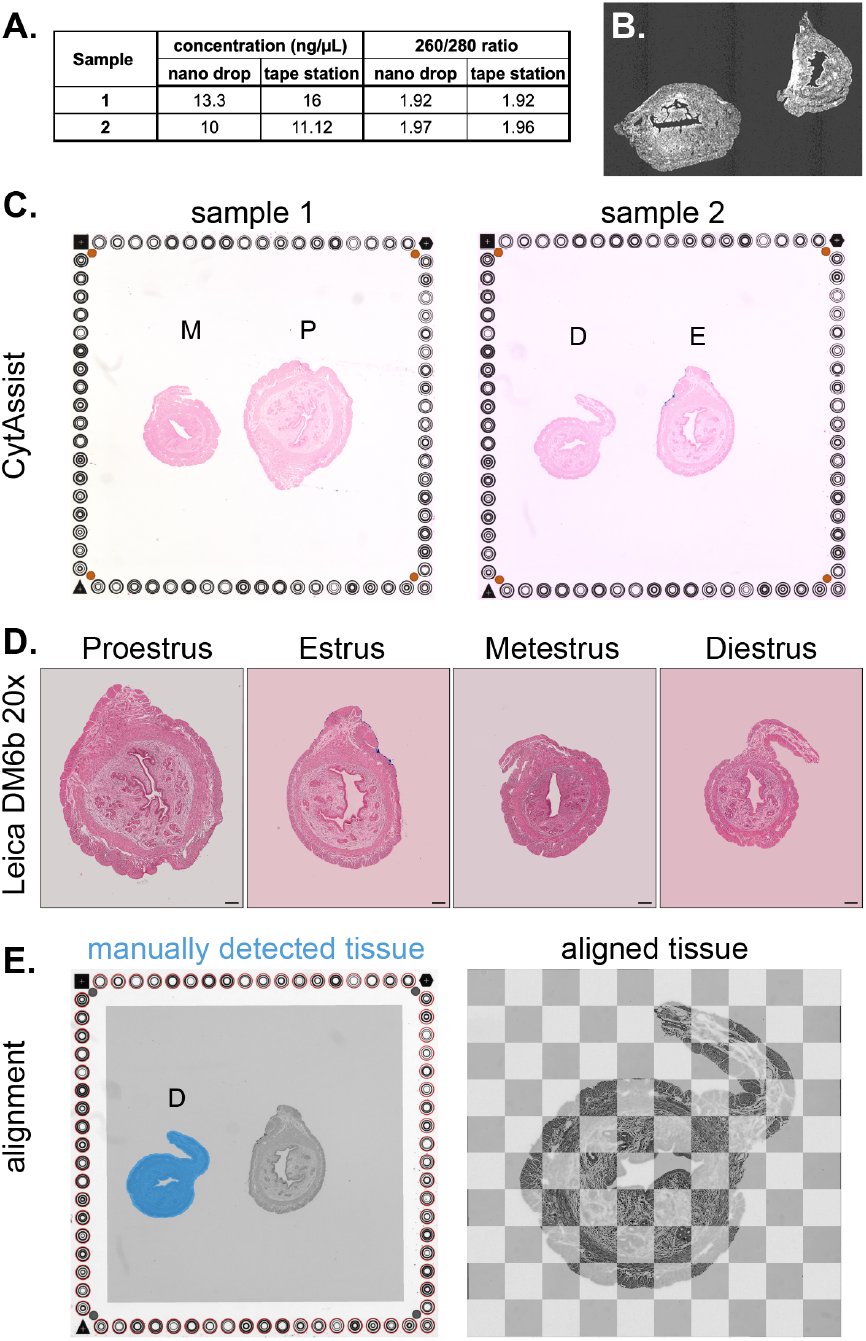
Quality control of uterine samples. A) RNA quality was assessed using nanodrop and Tapestation for each block. RNA was collected from 4 ribbons per block, and concentration (ng/µL) and 260/280 ratio was determined. B) RNA quality was assessed using control RNAscope probes. C) CytAssist images showing each sample and its associated uterine tissue. Sample 1 contained metestrus (M) and proestrus (P) tissue. Sample 2 contained diestrus (D) and estrus (E) tissue. D) Leica DM6b 20X images of each uterine tissue. Scale bar = 1mm. E) Alignment between the high magnification images from the Leica DM6b with low resolution images from the CytAssist. Tissue was manually detected (blue outline) and aligned tissue was visualized using a checkerboard pattern.

### Sequencing quality control

We performed quality control on our sequencing dataset. To achieve this, the total genes per spot were visualized on the section for each sample (**Figure 3A-D**). Luminal and glandular epithelium and the longitudinal smooth muscle layer displayed the highest number of genes per spot compared to other cell compartments like the stroma and circular smooth muscle layer (**Figure 3A-D**). The average number of Unique Molecular Identifiers (UMIs) per spot and genes per spot were calculated for each stage and visualized using violin plots (**Figure 3E-F**). We found similar average numbers of UMIs and genes per spot across each stage. Together, these quality control measures support that we have high-quality sequencing data.

**Figure 3.**
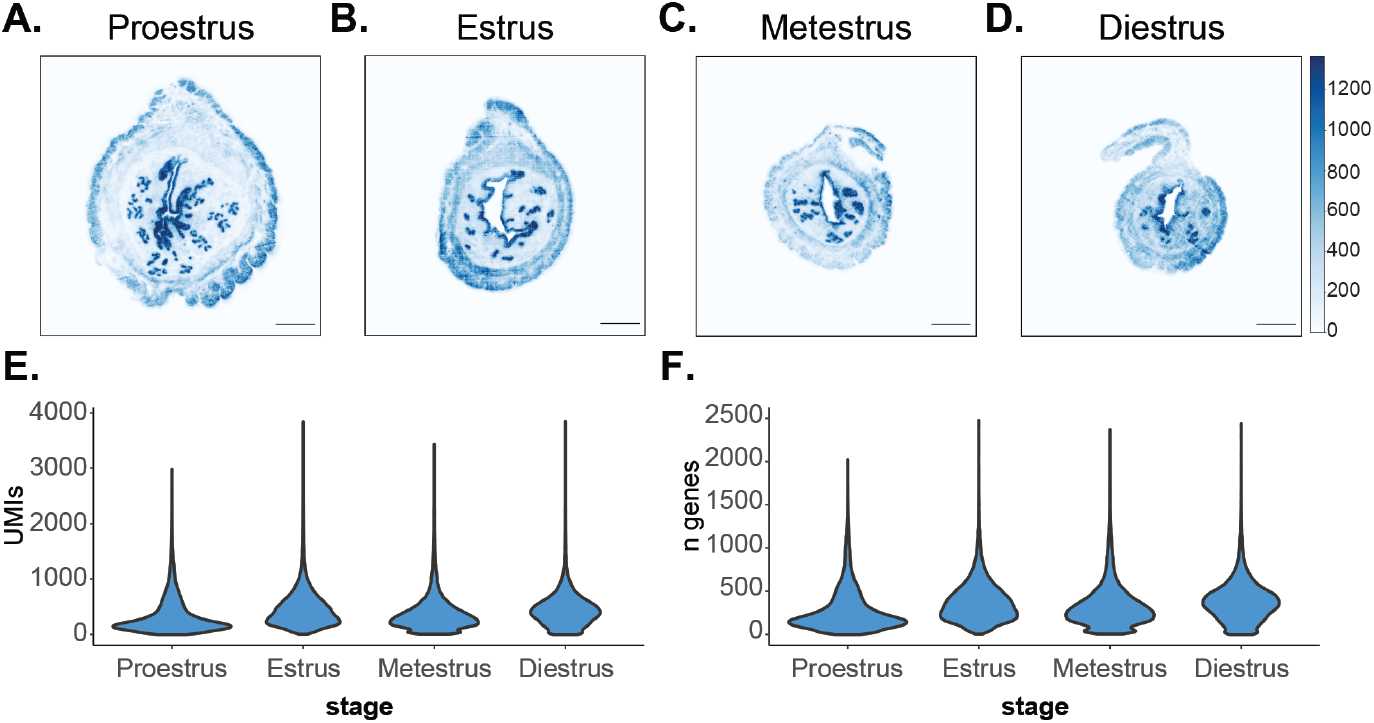
Technical validation of spatial datasets. Total RNA unique molecular identifier (UMI) counts per spot identified at: A) Proestrus, B) Estrus, C) Metestrus, and D) Diestrus, scale bar = 500µm throughout. Violin plots of the number of E) Unique Molecular Identifiers (UMIs) per cycle stage and F) genes identified per cycle stage.

### Integrated clustering and annotation of cell types

After quality control, we binned spots at 8 µm, clustered these spots based on transcriptional profile, and visualized clusters using UMAP embeddings (**Figure 4A**). We used a combination of marker genes, prior knowledge, and H&E spatial location to classify populations of stromal fibroblasts, smooth muscle, epithelium, immune cells, and endothelium. The following marker genes were used to classify the clusters: Longitudinal smooth muscle (LSM): *Tnnt2, Synm, Postn*; Outer circular smooth muscle (O-CSM): *Igfbp2, Acta1, Amhr2*; Inner circular smooth muscle (I-CSM): *Tpm2, Acta1, Adrb2*; Contractile smooth muscle (Contractile SM): *Smtn, Myh11, Tagln, Tpm2*; Luminal epithelium (LE): *Wnt7a, Wnt7b, Wnt11*; Glandular epithelium (GE): *Cxcl15, Foxa2*; Major stroma (M stroma): *Hsd11b2, Foxl2, Col6a4*; Peri-luminal epithelium stroma (P-LE stroma): *Col26a1, Col18a1, Wnt16*; Glandular epithelium/stroma mixture (GE-stroma mix): *Cldn10, Clu, Sprr2f*; Endothelium 1 – veins (Endo 1): *Pecam1, Emcn, Flt1, Ptprb*; Endothelium 2 – arteries (Endo 2): *Sema3g, Jag2, Hey2*; Endothelium 3 – proliferative (Endo 3): *Prc1, Foxm1, Top2a, Mki67, Pecam1, Eng*; Vascular smooth muscle (Vascular SM): *Notch3, Mgp*; Lymphatic endothelium 1 (Lymph Endo 1): *Prox1, Flt4, Lyve1*; Lymphatic endothelium 2 (Lymph Endo 2): *Prox1, Flt4*; Lymphatic endothelium 3 (Lymph Endo 3): *Prox1, Flt4, Lyve1*; Immune cell 1 - B cells, T cells (Immune 1): *Ms4a1, Cxcl9, Ighd, Cd79a, Cd4*; Immune cell 2 - antigen presenting cells (Immune 2): *C1qa, C1qb, H2-Ab1, H2-Aa, H2-Eb1*; Immune cell 3 - T cells, NK cells (Immune 3): *Il2rb, Ighm, Ptprc, Gzma, Klre1*; Mast cells: *Tpsab1, Cma1, Cpa3*. We also identified 3 additional clusters that remain unclassified based on either gene enrichment or spatial location (Unknown 1-3). The number of spots per cluster were plotted, showing that most cells were of the major stromal fibroblast population, followed by several smooth muscle clusters, and then epithelium (**Figure 4B**). In addition, the number of spots per cluster per cycle stage were visualized using a stacked bar plot (**Figure 4B**). These results showed that for some clusters, the number of spots per cluster varied with each cycle stage. For example, the I-CSM cluster has more spots at proestrus compared to other cycle stages (**Figure 4B, first column**). By contrast, Immune 1 (B and T cells) have more spots at estrus compared to other cycle stages (**Figure 4B, last column**). This immune infiltration is expected considering that uteri in estrus are at heightened inflammatory states^29^, and most immune populations are enriched in the uterus at estrus when assessed by flow cytometry^40^. Finally, the top three enriched genes per cluster, identified using the *FindMarkers* function in Seurat, were plotted as a heatmap (**Figure 4C**). These data suggest important transcriptional and spatial dynamics of myriad cell populations across the estrous cycle.

**Figure 4.**
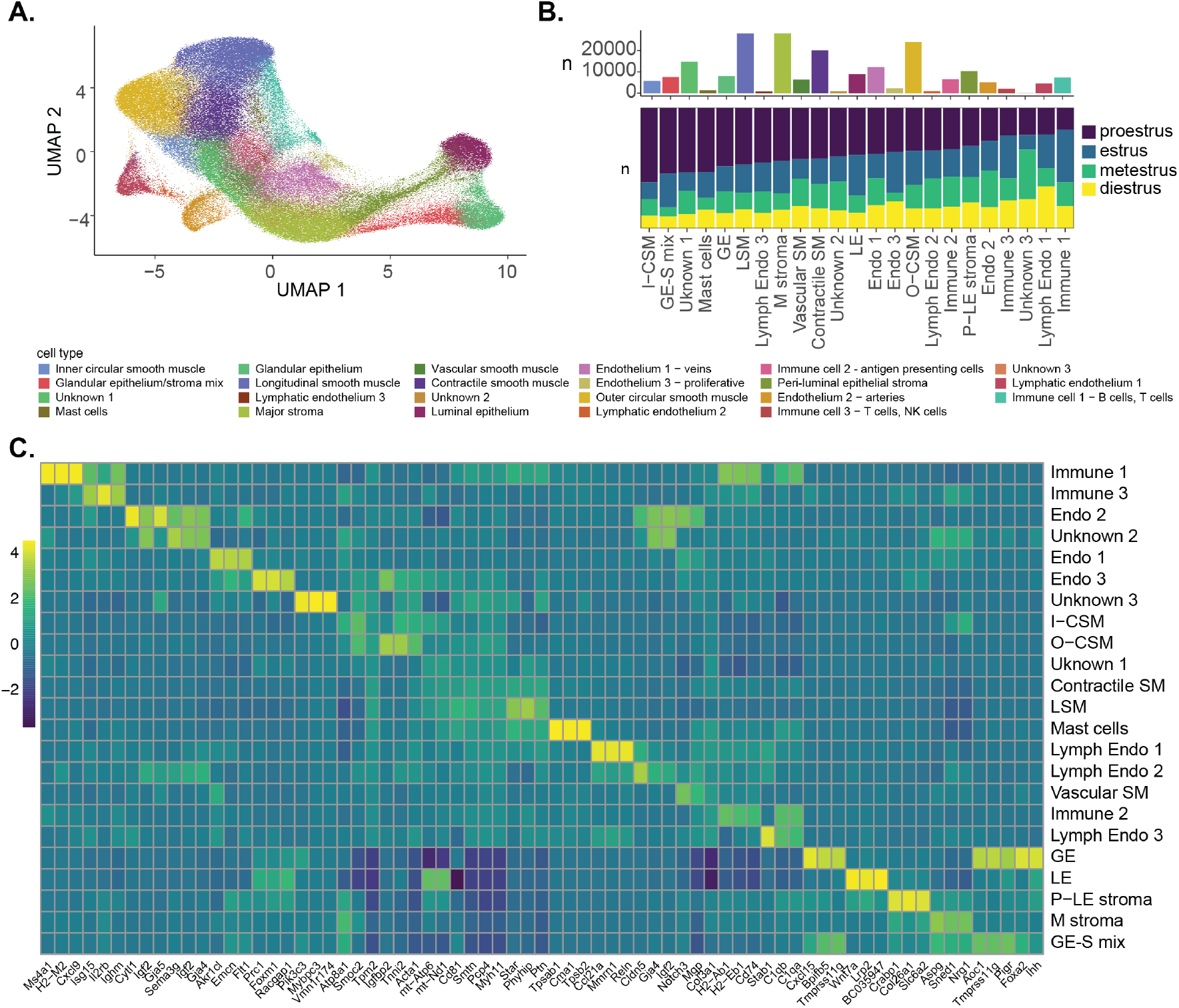
Integration of cell clusters across the estrous cycle. A) UMAP embedding displaying cell clusters integrated across the estrous cycle. B) Combination bar plot showing total number of 8 µm bins per cluster, and stacked bar plot displaying relative proportion of 8 µm bins per cluster per cycle stage. Bar plot is color coded by cell type (see legend to the left). The same order of clusters is displayed in the stacked bar plot and is color coded by cycle stage, where purple = proestrus, blue = estrus, green = metestrus, and yellow = diestrus. C) Heatmap displaying top 3 most enriched genes expressed in each cluster. Values represent the sum of all spots assigned to a given cell type across all four samples.

### Spatial visualization of gene expression and cell types

Each cluster was precisely mapped back onto the H&E images (**Figure 5A-D**). The inset for these images zooms onto glandular epithelium (teal) surrounded by glandular epithelium/stromal mixture (bright red), and other surrounding populations. We were able to use spatial distribution of smoothed 2 µm expression data (see methods) to confirm cell-specific gene expression with previously published literature. For example, *Foxa2* and *Cxcl15* are specifically expressed by glandular epithelium (**Figure 5E-F**)^41,42^. *Ihh* is expressed by both luminal epithelium and gland epithelium (**Figure 5G**)^43^. *Wnt7a* is expressed specifically by the luminal epithelium (**Figure 5H**)^44^. Finally, to highlight the increased resolution achieved by smoothing the 2 µm expression data, we zoomed in on gland epithelium at proestrus (**Figure 5I**) and compared *Foxa2* expression binned into 8 µm squares (**Figure 5J**) to either raw or smoothed 2 µm expression profiles (**Figure 5K, L** respectively). This analysis serves as validation that our dataset is high enough quality to identify discrete cell types within the uterus. Thus, we expect these data to serve as a valuable resource that will enable discoveries about disease genes and their expression patterns in the myriad cell types of the cycling uterus.

**Figure 5.**
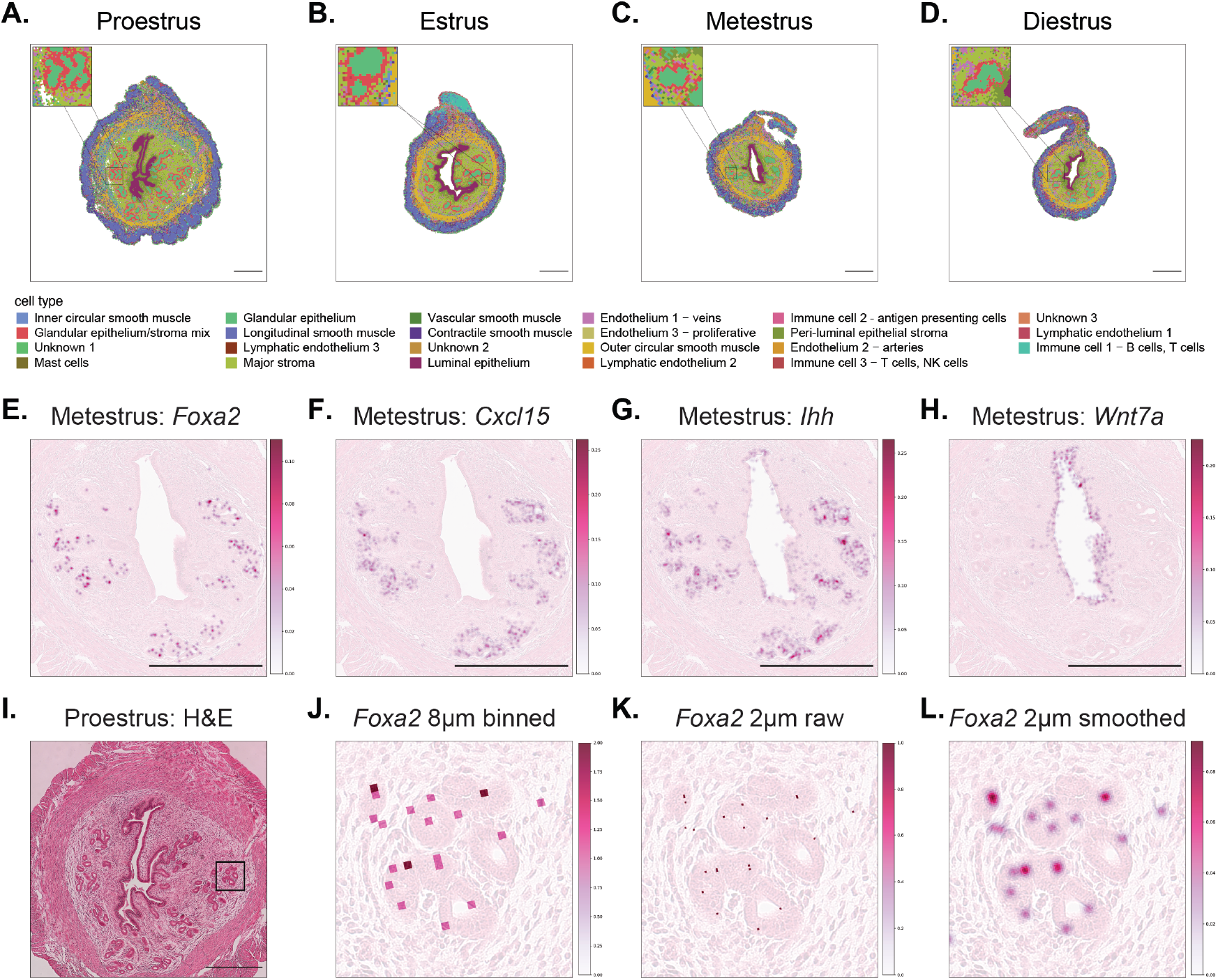
Annotation of cell clusters across the estrous cycle. Integrated clusters mapped onto estrous samples, where inset shows gland epithelium and associated stromal populations at A) proestrus, B) estrus, C) metestrus, and D) diestrus. E) Smoothed 2 µm *Foxa2* expression in glandular epithelium at metestrus. F) Smoothed 2 µm *Cxcl15* expression in glandular epithelium at metestrus. G) Smoothed 2 µm *Ihh* expression in luminal and glandular epithelium at metestrus. H) Smoothed 2 µm *Wnt7a* expression in luminal epithelium at metestrus. I) H&E image of the proestrus sample. The box denotes the zoomed-in area for J-L. J) *Foxa2* expression counts binned into 8 µm squares. K) *Foxa*2 expression displayed as raw 2 µm counts. L) *Foxa2* expression displayed as 2 µm data smoothed using kernel density estimation (see methods). Scale bar = 500µm throughout.

## Potential limitations of the dataset

While our dataset has many strengths, it is important to note some of the potential limitations. First, our dataset represents single-cell-<u>scale</u> transcriptomes, not true single cells, therefore classifications may be a combination of cell types rather than individual cell types. This is most readily depicted by the glandular epithelium/stroma mixture cluster, where the heatmap (**Figure 4C**) shows a combination of two other clusters’, glandular epithelium and major stroma, marker gene expression. Second, we used a probe-based Visium HD slide which means that the slide does not have 100% coverage of the genome. Instead, expression of ∼19,059 genes are targeted by probes. Therefore, we may be missing some genes from our dataset.

## Acknowledgements

This study was supported by start-up funds from the CU Anschutz Medical Center, School of Medicine, Department of Pediatrics, Section of Developmental Biology to E.C.R. This study was supported in part by the National Institutes of Health P30CA069345 funded Research Histology (RRID: SCR_021994), Human Immune Monitoring (RRID: SCR_021985), and Genomics Shared Resources (RRID: 021984). The authors would also like to acknowledge the vivarium staff for their support maintaining the mouse colony, and the McKey Lab at CU Anschutz Medical Center for their support and enthusiasm regarding this spatial transcriptomics dataset.

## Author contributions

E.U. designed the study, collected and prepared samples for analysis, and assisted in drafting the manuscript. T.J.G. performed the bioinformatic analyses and assisted in drafting the manuscript. L.J.B. assisted in designing the study and sample collection. E.C.R. designed the study and drafted the manuscript.

